# 3D reconstruction of SARS-CoV-2 infection in ferrets emphasizes focal infection pattern in the upper respiratory tract

**DOI:** 10.1101/2020.10.17.339051

**Authors:** Luca M. Zaeck, David Scheibner, Julia Sehl, Martin Müller, Donata Hoffmann, Martin Beer, Elsayed M. Abdelwhab, Thomas C. Mettenleiter, Angele Breithaupt, Stefan Finke

## Abstract

The visualization of viral pathogens in infected tissues is an invaluable tool to understand spatial virus distribution, localization, and cell tropism *in vivo*. Commonly, virus-infected tissues are analyzed using conventional immunohistochemistry in paraffin-embedded thin sections. Here, we demonstrate the utility of volumetric three-dimensional (3D) immunofluorescence imaging using tissue optical clearing and light sheet microscopy to investigate host-pathogen interactions of pandemic SARS-CoV-2 in ferrets at a mesoscopic scale. The superior spatial context of large, intact samples (> 150 mm^3^) allowed detailed quantification of interrelated parameters like focus-to-focus distance or SARS-CoV-2-infected area, facilitating an in-depth description of SARS-CoV-2 infection foci. Accordingly, we could confirm a preferential infection of the ferret upper respiratory tract by SARS-CoV-2 and emphasize a distinct focal infection pattern in nasal turbinates. Conclusively, we present a proof-of-concept study for investigating critically important respiratory pathogens in their spatial tissue morphology and demonstrate the first specific 3D visualization of SARS-CoV-2 infection.

## Introduction

In December 2019, a novel coronavirus (2019-nCoV) associated with viral pneumonia emerged in Wuhan, Hubei Province, China [1–4]. The virus was subsequently designated as severe acute respiratory syndrome coronavirus 2 (SARS-CoV-2) [5] and identified to be the causative agent of COVID-19 (Coronavirus disease 2019). Patients most commonly present with fever, cough, fatigue, and dyspnea [6–9]. About one out of five patients develops severe disease [10,11]. The outbreak was declared a “public health emergency of international concern” on January 30, 2020, and a pandemic on March 11, 2020. As of October 12, 2020, 37,109,851 confirmed cases and 1,070,355 confirmed death were reported in 235 countries [12].

SARS-CoV-2 is an enveloped virus with a single-stranded RNA genome of positive polarity and has been classified as a member of the *Coronaviridae* family, genus *Betacoronavirus* [5]. Including SARS-CoV-2, there are currently two alpha-and five betacoronaviruses associated with human disease [13]. While most result in only mild respiratory illness, the three zoonotic betacoronaviruses SARS-CoV [14], SARS-CoV-2 [1–4], and MERS-CoV [15] (Middle East respiratory syndrome coronavirus) can cause severe respiratory disease. Bats presumably serve as natural reservoir for both SARS-CoV [16,17] and MERS-CoV [18–20], whereas palm civets [21] and dromedary camels [22] have been identified as the intermediate hosts for animal-human transmission for SARS-CoV and MERS-CoV, respectively. Viruses closely related to SARS-CoV-2 have been found in bats [2] and Malayan pangolins [23,24] but no direct transmission event or intermediate host species have been identified thus far.

Over the course of the current pandemic, tremendous research efforts have been undertaken to study the virus and its disease. Consequently, critical information on the virus, e.g. receptor usage [25], and necessary research tools, including reverse genetics systems [26,27], became rapidly available. Furthermore, a variety of animal studies to investigate susceptibility and suitability as animal models have been conducted in a number of animal species (reviewed in [28]): ferrets [29–31], hamsters [32–34], cats [31,35], dogs [31], raccoon dogs [36], rabbits [37], transgenic mice [38–40], pigs [30,31], cattle [41], monkeys [42–44], poultry [30,31,45], and fruit bats [30].

Within the respiratory tract, detection of viral antigen and RNA suggested a preferential replication of SARS-CoV-2 in the upper respiratory tract (URT) of ferrets [29–31], whereas viral antigen was detected in both the URT und lower respiratory tract (LRT) of Syrian hamsters [32,33]. In humans and non-human primates (NHPs), viral antigen detection indicates virus replication in both the URT and LRT [42,46,47].

Thus far, almost all approaches to detect and image SARS-CoV-2 infection in tissues have been based on conventional immunohistochemistry (IHC) of paraffin-embedded thin sections. However, by omitting the spatial context, thin tissue sections of only several micrometers in thickness bear the risk of incomplete or inaccurate description, particularly for focal infections. Recent developments in the field of tissue optical clearing (TOC) have facilitated the preservation of large intact tissue structures by turning them optically transparent. This eliminates the need of physical sectioning and allows acquisition of intact three-dimensional (3D) structures using only optical sectioning, e.g. in light sheet fluorescence microscopy (LSFM) (reviewed in [48]). Lately, the opportunities and advantages of TOC for virus research have been demonstrated in several studies [49–54]. While two approaches to 3D imaging of SARS-CoV-2-infected lung tissue have been described recently [55,56], neither of them is capable of direct visualization of SARS-CoV-2 infection via virus-specific antigen staining.

In our study, we provide a first complete 3D overview of SARS-CoV-2 infection in the ferret model. By staining for the viral nucleocapsid protein (SARS-CoV-2 N), we were able to directly visualize and localize SARS-CoV-2-infected foci within large volumes of the ferret respiratory tract. Direct visualization further allowed detailed description of the foci in their spatial context. To the best of our knowledge, this is the first report of specific 3D reconstruction of SARS-CoV-2 infection as well as the first report of 3D visualization of respiratory virus infection in nasal turbinates using LSFM.

## Materials and Methods

### Cells and viruses

VeroE6 cells (CCLV-RIE 0929, Collection of Cell Lines in Veterinary Medicine [CCLV], Friedrich-Loeffler-Institut, Germany) were maintained in Minimal Essential Medium (Gibco, USA) supplemented with 10% fetal bovine serum (Biowest, France) and non-essential amino acids (Gibco).

SARS-CoV-2 isolate 2019_nCoV Muc-IMB-1 (kindly provided by Roman Wölfel, German Armed Forces Institute of Microbiology, Germany) was propagated on VeroE6 cells. The complete sequence is available through GISAID under the accession number ID_EPI_ISL_406862 and name “hCoV-19/Germany/BavPat1/2020”.

### Antibodies and chemical reagents

For the detection of SARS-CoV-2 infection, a 1:1 mixture of hybridoma cell culture supernatants of anti-SARS-CoV-1 N mouse monoclonal antibody clones 4E10A3A1 (RRID:AB_2833160) and 4F3C4 (RRID:AB_2833162) [57] at a dilution of 1:5 or a polyclonal rabbit anti-SARS-CoV-1 N (RRID:AB_838838; Novus Biologicals, USA) at a dilution of 1:250 were used. Alexa Fluor™ 488/568/647-conjugated antibodies against mouse IgG and rabbit IgG were used as secondary antibodies (1:500; Invitrogen, USA).

A detailed list of the chemicals and reagents used for the immunostaining and optical clearing of SARS-CoV-2-infected tissue samples, and their suppliers is provided in Supplementary Table S1.

### Virus infection and immunofluorescence staining of mammalian cell cultures

5 × 10^5^ VeroE6 cells were seeded on coverslips one day prior to infection with 1 × 10^6^ TCID_50_ of SARS-CoV-2 isolate 2019_nCoV Muc-IMB-1. Infected VeroE6 cells were then fixed 24 h post-infection with 4% paraformaldehyde (PFA) for 20 min. Following permeabilization with 0.5% Triton X-100/PBS for 15 min, cells were blocked with 10% normal donkey serum in 0.1% Tween-20/PBS (PBS-T) for 30 min. Primary antibodies against SARS-CoV N were applied for 1 h at room temperature in 1% normal donkey serum/PBS-T, followed by three washes with PBS and incubation with the secondary antibody for 1 h at room temperature in 1% normal donkey serum/PBS-T. Nuclei were counterstained with Hoechst33342 (Invitrogen) and samples were embedded in ProLong™ Glass AntiFade Mountant (Invitrogen) for analysis by confocal laser-scanning microscopy.

### Tissue samples of SARS-CoV-2-infected ferrets

In a previous study on experimental transmission of SARS-CoV-2 among different animal species, ferrets were inoculated intranasally with 105 TCID50 of SARS-CoV-2 isolate 2019_nCoV Muc-IMB-1 [30]. Tissues were collected in 10% neutral-buffered formalin and fixed for at least 21 days to ensure complete virus inactivation. In this study, nasal conchae, trachea, and lung tissue samples from infected ferrets euthanized on day 4 post-infection were analyzed.

### Immunofluorescence staining of high-volume tissue sections

Large sections of respiratory tissues (≥ 150 mm^3^) were immunostained according to a modified iDISCO protocol [50,58]. All incubation steps were conducted with slight agitation and, if not indicated otherwise, at room temperature.

To this end, formaldehyde-fixed tissues were washed three times for at least 1 h each in PBS. Nasal conchae were furthermore decalcified for 4-7 days in Formical-2000™ (Statlab, USA). Samples were trimmed to the sizes and volumes described above, and bleached overnight in 5% H2O2 in PBS at 4 °C. For permeabilization, the tissue samples were first incubated twice for 3 h each with 0.2%Triton X-100/PBS at 37 °C and subsequently in 0.2% Triton X-100/20% DMSO/0.3 M glycine/PBS for 2 days at 37 °C. Following a blocking step with 6% normal donkey serum/0.2% Triton X-100/10% DMSO/PBS for 2 days at 37 °C, primary antibodies were diluted in 3% normal donkey serum/5% DMSO in PTwH (0.2% Tween-20 in PBS with 10 µg/mL heparin) and applied for 4 days at 37 °C. Unbound antibody was removed by washing the samples 4-5 times over the course of a day, leaving the final wash on overnight. Secondary antibodies were diluted in 3% normal donkey serum/PTwH and the samples were incubated for another 4 days at 37 °C. Washing was performed as described for the primary antibody.

### Ethyl cinnamate (ECi)-based tissue optical clearing

Immunostained tissue sections were cleared with an adjusted ECi-based protocol [59]. All incubation steps were conducted with slight agitation.

The samples were dehydrated in a graded ethanol series (30% [v/v], 50%, 70%, and twice in 100%; each for ≥ 8 h at 4 °C, diluted in aqua ad iniectabilia, and pH-adjusted to 9). Following a two-hour wash with *n*-hexane at room temperature [60], *n*-hexane was gradually replaced with the clearing agent ECi and samples were incubated until optically transparent.

### Light sheet microscopy of optically clear tissue samples

Light sheet micrographs of optically clear and immunostained respiratory tissues from SARS-CoV-2-infected ferrets were acquired with a LaVision BioTec Ultramicroscope II (LaVision, Germany). The microscope was equipped with an Olympus MVX-10 zoom body (magnification range: 0.63x – 6.3x, total magnification: 1.26x – 12.6x; Olympus, Japan), an Olympus MVPLAPO 2x objective (NA = 0.5), a LaVision laser module with four laser lines (488 nm, 561 nm, 639 nm, and 785 nm), and a Andor Zyla 5.5 sCMOS Camera (Andor Technology, UK) with a pixel size of 6.5 µm2. To visualize tissue morphology, non-specific autofluorescence was excited with the 488 nm laser. Excitation lines 561 nm and 639 nm were used to excite Alexa Fluor™ 568 and Alexa Fluor™ 647, respectively. Channels of a high-volume 3D image were acquired sequentially with a z-step size of 2 µm, a light sheet width of 100%, and a light sheet thickness of 3.89 µm (NA = 0.156). Acquisition was done with ImSpector (v7.0.124.0).

### Confocal laser-scanning microscopy (CLSM)

Confocal images were acquired with a Leica DMI6000 TCS SP5 confocal laser-scanning microscope (Leica Microsystems, Germany) equipped with a 63x/1.40 oil immersion HCX PL APO objective and a 40x/1.10 water immersion HC PL APO objective. Fluorescence was recorded sequentially between lines with a pinhole diameter of 1 Airy unit and z-step sizes of 0.35 µm. Acquisition was done with LAS AF (v2.7.3.9723).

For high-resolution confocal laser-scanning analysis of cleared and immunostained tissue samples, they were sectioned into 1-mm-thick slices using a stainless steel tissue matrix (World Precision Instruments, UK). Tissue slices were then mounted in 3D-printed imaging containers as described before [50]. Tissue morphology was reconstructed from non-specific tissue autofluorescence via excitation with a 405 nm UV laser diode.

### Image processing and analysis

Image visualization and analysis were performed with arivis Vision4D (v3.2). If necessary, channels were background corrected. CLSM-acquired image stacks of subsectioned volumetric tissue samples were denoised. To quantify relations between SARS-CoV-2 infection foci, they were segmented. The shortest distances between foci were measured using the segment operation “Distances”. To calculate the area of SARS-CoV-2-infected tissue, surface areas of the segmented objects were extracted and divided by two to account only for the surface of the object facing outwards. Lookup tables of multicolor images were selected for maximum accessibility.

## Results

### LSFM provides a unique insight into the spatial distribution of SARS-CoV-2 in intact nasal turbinates

By combining LSFM with optically cleared samples of the ferret respiratory tract (Figure 1A), we aimed to shed light on the infection environment and spatial context of SARS-CoV-2 infection. A commercially available polyclonal serum (designated as #1), which has been used for SARS-CoV-2 detection by conventional IHC [30,42], and a mix of two monoclonal antibodies (designated as #2) against SARS-CoV N were tested on virus-infected VeroE6 cells and confirmed to be cross-reactive with SARS-CoV-2 N (Figure 1B). Following immunostaining, ferret tissue samples, including the partly ossified nasal conchae, were successfully turned optically transparent using a recently established ethyl cinnamate (ECi)-based approach [59] (Figure 1C).

**Figure 1:**
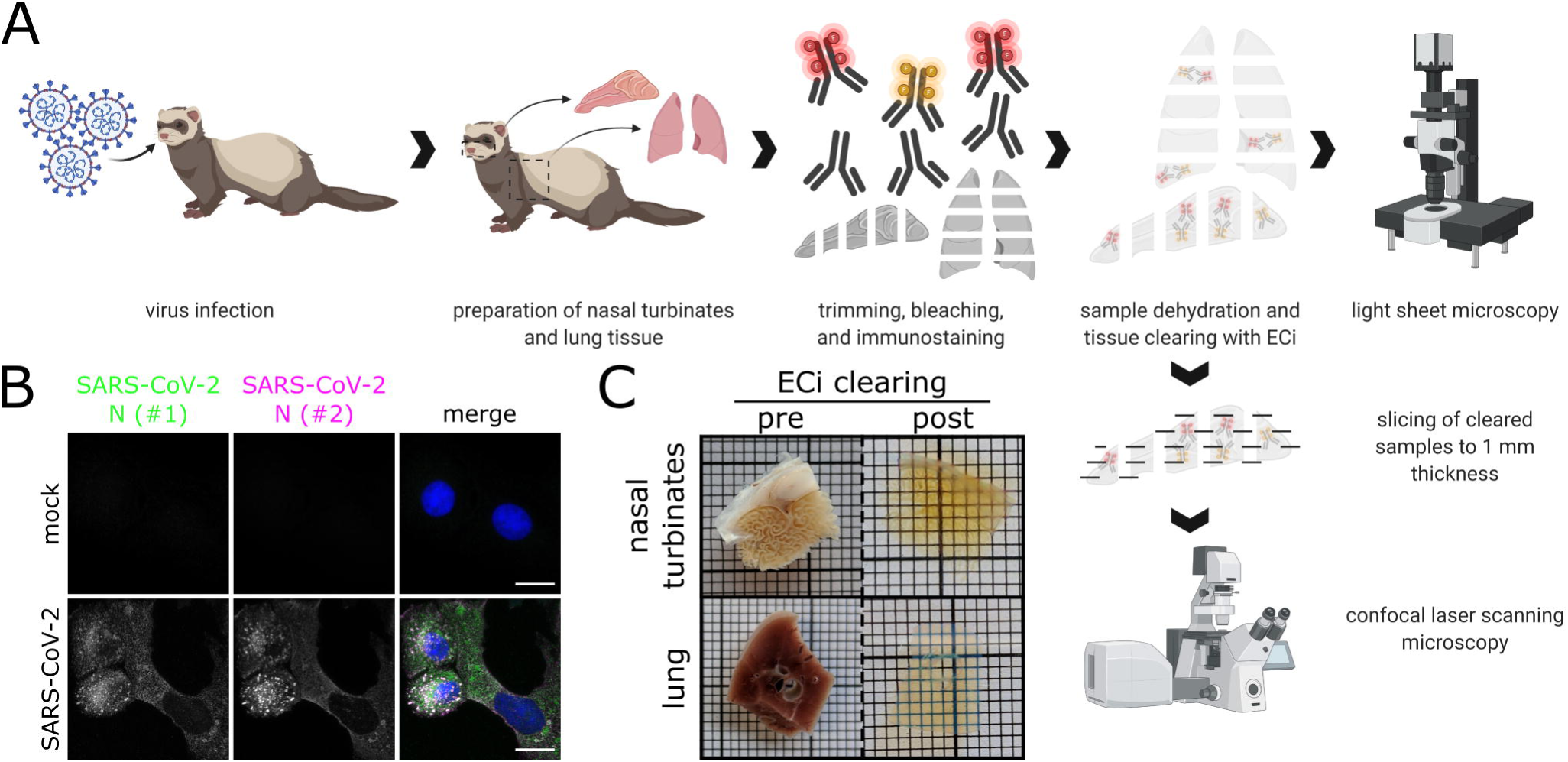
Workflow for correlative LSFM-CLSM of SARS-CoV-2-infected ferret tissues. **(A)** Nasal conchae and lungs tissue from SARS-CoV-2-infected ferrets were collected, trimmed, and immunostained against SARS-CoV-2 N protein. Fully dehydrated and optically transparent samples were acquired *in toto* with a light sheet microscope and subsequently subsectioned to 1 mm-thick sections for correlative confocal laser-scanning microscopy. **(B)** Representative immunostaining for SARS-CoV-2 N in infected VeroE6 cells using a commercially available polyclonal anti-SARS-CoV N serum (#1, green) and a monoclonal anti-SARS-CoV N mix (#2, magenta) confirms antibody specificity. Blue: Hoechst33342. Scale bars = 15 µm. **(C)** Representative ferret respiratory tract samples before (left) and after (right) immunostaining and ECi-based optical clearing. The photographs from the lung sections (bottom) show two independent samples. Edge length of grid square: 1 mm.

Full translucency of ferret nasal turbinates enabled LSFM acquisition of a > 200 mm3 (6.69 gigavoxels per channel; ∑ = 20.07 gigavoxels)-sized tissue sample (Figure 2A and Supplementary Movie S1). While there were some unspecific signals detectable in the SARS-CoV-2 N-stained sample (individual green or magenta spots), they could be clearly distinguished from specific SARS-CoV-2 detection by the absence of colocalization (white) of the signals from either antibody (Figure 2B and Figure S1). Within the about 4-mm-thick URT sample (4 days post-infection), multiple comparatively small SARS-CoV-2 infection hot spots were visualized (Figure 2B). They were detected in both the *Concha nasalis dorsalis* (Figure 2B, ROIs [region of interests] 1 & 2) and the *Concha nasalis ventralis* (Figure 2B, ROI 3). Overall, these data provide the proof of concept for the feasibility of TOC-assisted LSFM analysis for SARS-CoV-2.

**Figure 2:**
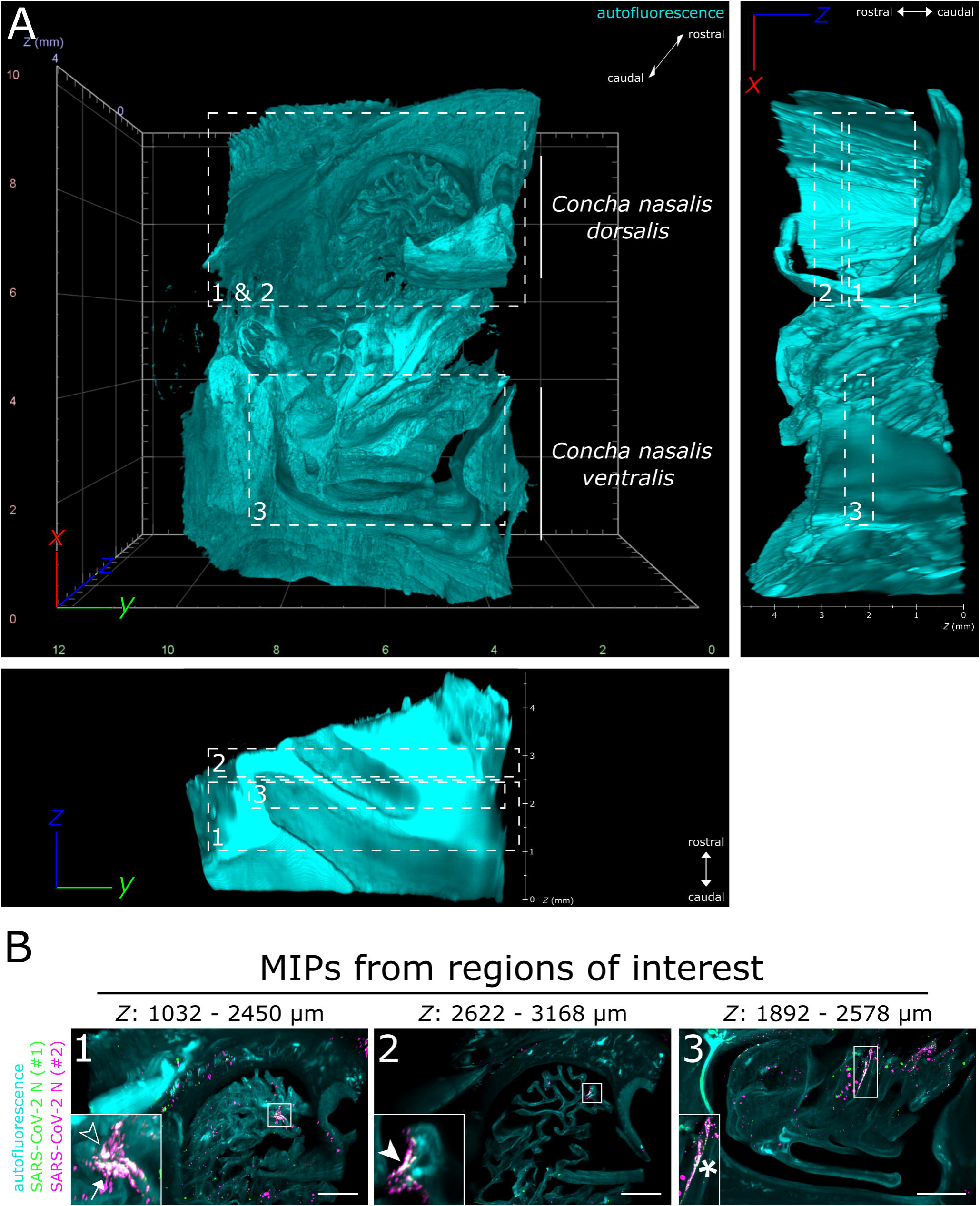
LSFM is able to visualize SARS-CoV-2 infection in nasal turbinates within a high spatial context. **(A)** The tissue structure of the nasal conchae (> 200 mm3; 4 days post-infection) was reconstructed using tissue autofluorescence (cyan) and is depicted as volumetric projection from three viewing angles. Anatomical terms of location are provided for orientation. Edge length of grid squares = 2 mm. Total magnification = 1.26x. **(B)** Maximum intensity projections of the regions of interest (1-3) highlighted in (A). SARS-CoV-2 infection is characterized by colocalization of both SARS-CoV-2 N stainings (#1, green; #2, magenta) and results in white coloring (inset). Four distinct SARS-CoV-2 infection foci are highlighted (filled-in arrowhead [A1], outlined arrowhead [A5], arrow [A7], and asterisk). Foci will hereafter be referred to via their respective indicator or designation in square brackets. Ranges of the MIPs in the *z*-dimension are provided above the respective image. MIP = maximum intensity projection. Scale bar = 1 mm.

### SARS-CoV-2 infection in the upper respiratory tract of ferrets is characterized by an oligofocal infection pattern

To achieve a more in-depth analysis of the individual SARS-CoV-2 infection foci, LSFM image stacks of infected areas were acquired using a higher magnification (total magnification of 8x for Figure 3 vs. 1.26x for Figure 2), thus, increasing image resolution while maintaining the complete spatial context (Figure 3).

**Figure 3:**
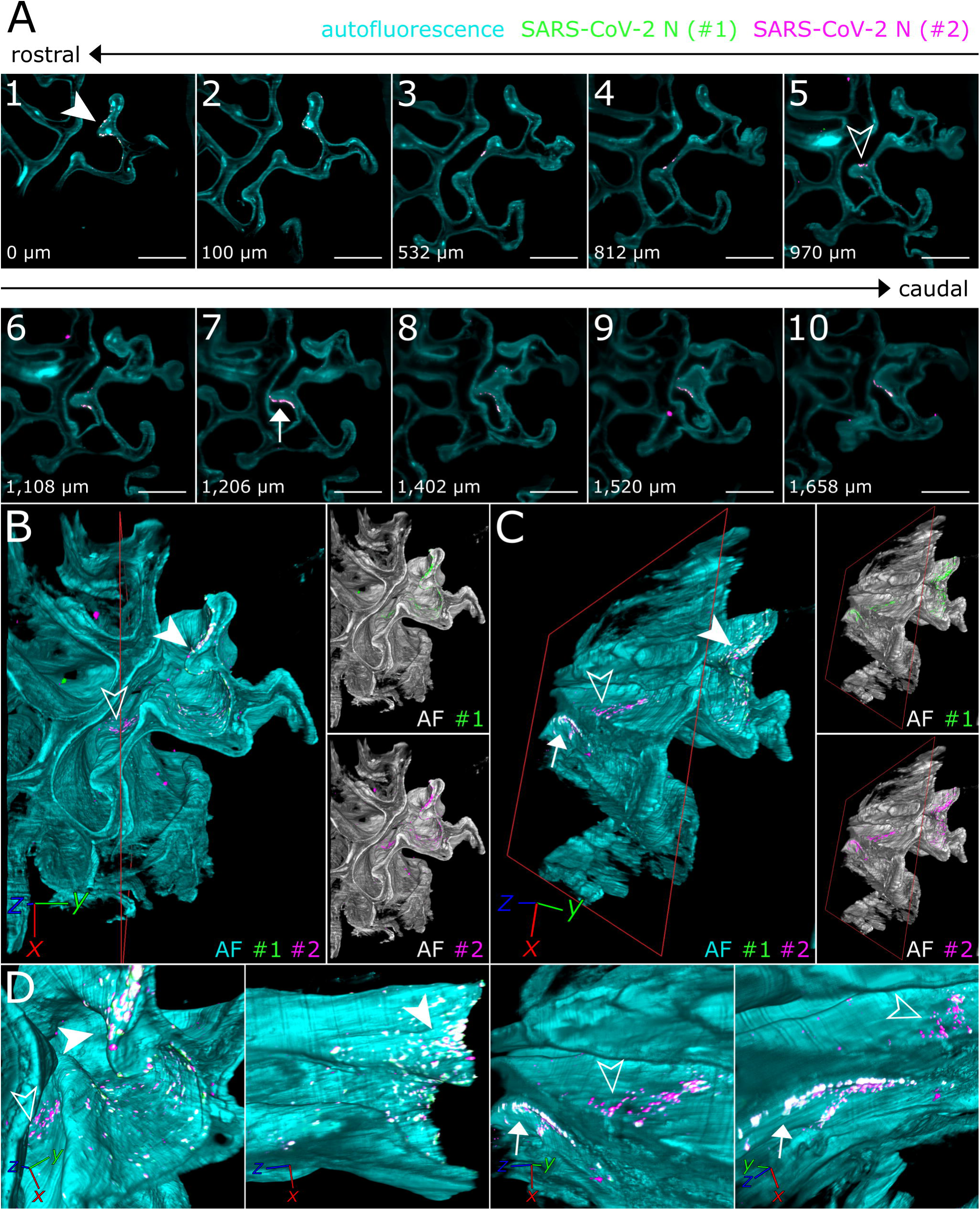
3D detail views highlight oligofocal SARS-CoV-2 infection pattern in nasal turbinates at 4 days post-infection. **(A)** Tomographic representation of three individual SARS-CoV-2 foci (filled-in arrowhead [A1], outlined arrowhead [A5], and arrow [A7]) from ROIs 1 & 2 in Figure 2 along a length of 1,658 µm. The relative distance to plane #1 is indicated in the bottom left corner. Cyan = autofluorescence; green = SARS-CoV-2 N #1; magenta = SARS-CoV-2 N #2. Scale bar = 400 µm. Total magnification = 8x. **(B,C)** Volumetric projections of the detail view from two angles. Clipping at the indicated plane (red) reveals the third SARS-CoV-2 foci (C, arrow), which is hidden behind nasal turbinate tissue in (B). Single channel views further emphasize the colocalizing pattern of both SARS-CoV-2 N stainings (#1, green; #2, magenta). Cyan/grayscale = autofluorescence (AF). **(D)** Close-ups of the three individual infection foci. The angle of the respective image is indicated in the bottom left corner.

Virtually traveling through an image stack of the ROIs 1 & 2 from Figure 2, which was acquired accordingly, corroborated the presence of three individual, well delimitable, and distinguishable SARS-CoV-2 infection foci (Figure 3A, images 1 [filled-in arrowhead], 5 [outlined arrowhead], and 7 [arrow]). A volumetric reconstruction of this image stack was able to convey spatial relationships between the individual infection spots (Figure 3B-D). Specificity of SARS-CoV-2 N detection was again confirmed by colocalization of both independent antibody stainings (Figure 3B and C, right side). Consequently, linear distances between foci and foci areas were quantified (Figure 4 and Table 1). This increased spatial context within the nasal turbinate sample is further reinforced by the fact that the SARS-CoV-2 infection focus from Figure 3A7 (arrow) is buried deeper within the sample and cannot be seen from the frontal angle in Figure 3B. However, either by virtually clipping the sample as indicated (Figure 3B and C, red square) or using 3D rendering to virtually “fly through the sample” (Supplementary Movie S2), the infection site can be visualized. Taken together, this emphasizes the system’s flexibility to switch from broad, mesoscopic overviews to detailed, resolved close-ups. By maintaining the full infection environment, we were able to establish quantifiable relations between the individual SARS-CoV-2 foci and highlight the oligofocal infection pattern of SARS-CoV-2 in the URT of ferrets.

**Table 1:** Direct linear distances between, areas affected by, and volumes of segmented SARS-CoV-2 infection foci. Linear distances were calculated either as the distance between the center of two foci or as the shortest possible distance between the edges of two foci. The area affected by SARS-CoV-2 infection was measured by calculating the surface area of the segmented objects and dividing the resultant value by two, thus, only accounting for the surface facing outwards.

| linear distance [$\mu\text{m}$] | | A1 | A5 | A7 |
| --- | --- | --- | --- | --- |
| from the center | A1 | 0 | 777.4 | 1303.3 |
|  | A5 | 777.4 | 0 | 554.3 |
|  | A7 | 1303.3 | 554.3 | 0 |
| from the edge | A1 | 0 | 421.1 | 997.5 |
|  | A5 | 421.1 | 0 | 75.5 |
|  | A7 | 997.5 | 75.5 | 0 |
| area [$\mu\text{m}^2$] | | 90,212 | 75,389 | 181,431 |
| volume [$\mu\text{m}^3$] | | 465,818 | 443,207 | 1,533,316 |

**Figure 4:**
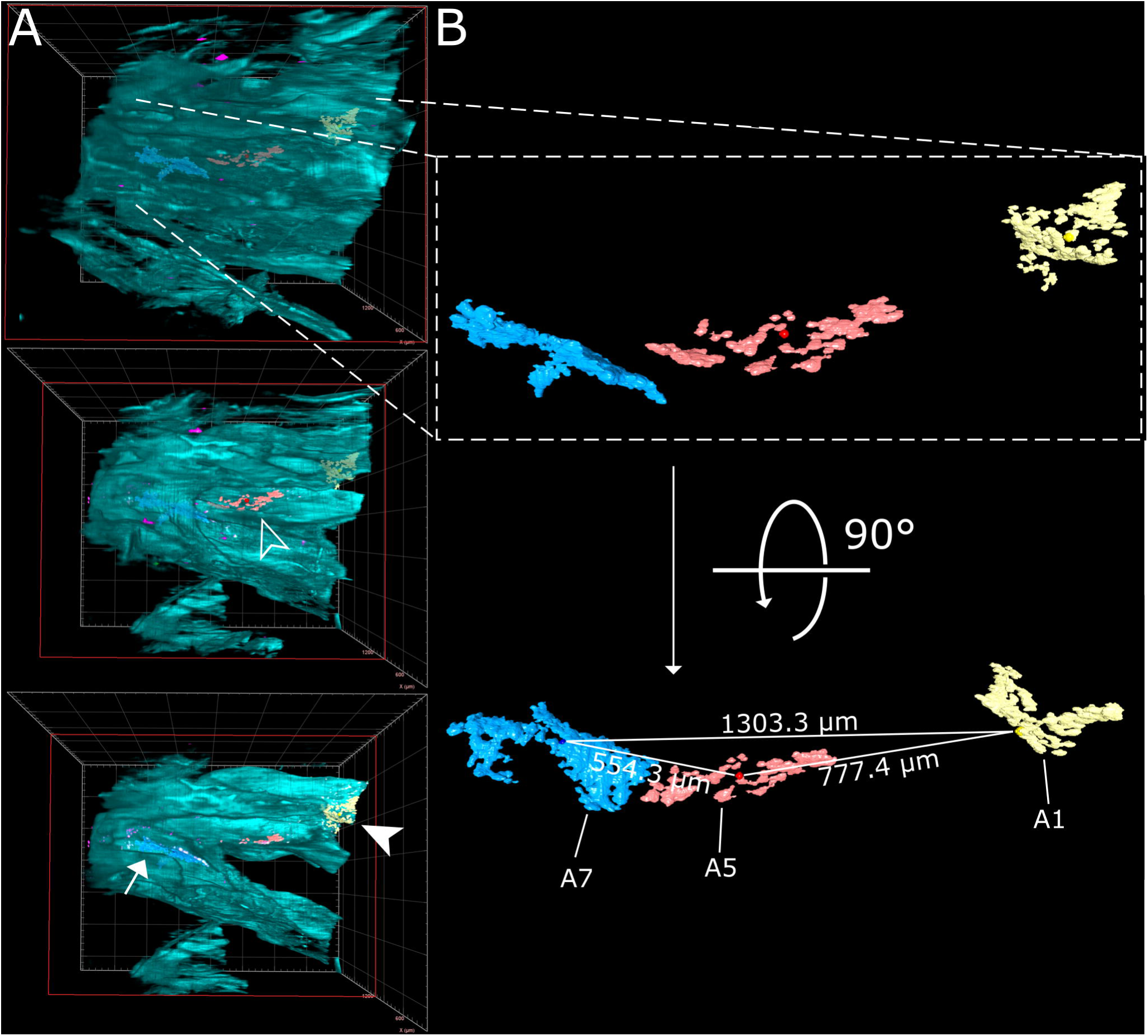
Virtual segmentation of SARS-CoV-2 infection foci at 4 days post-infection. **(A)** xz-view of the magnified nasal turbinate view from Figure 3, clipped at the indicated plane (red square). Segmented SARS-CoV-2 infection foci (A1: yellow, A5: red; A7: light blue) are visible through the autofluorescence reconstruction of the tissue morphology (cyan). Once they are uncovered by the clipping plane, they are highlighted with their respective indicator. **(B)** Detail and alternate viewing angle of segmented infection foci. A slightly darker sphere represents the respective foci center. The direct linear distances between the centers of each foci (from Table 1) are highlighted.

### CLSM acquisition of correlated regions of interest at subcellular resolution – infection of ciliated and non-ciliated cells in the nasal epithelium

While LSFM is ideally suited to generate a mesoscopic overview to analyze, for example, large-scale spatial virus distribution within virus-infected tissues, simultaneous resolution of subcellular details is not possible. Thus, following LSFM acquisition, we subsectioned the optically cleared high-volume tissue sample to 1-mm thick slices using a tissue matrix to achieve compatibility with the limited free working distances of CLSM objectives (Figure 1A).

Using the spatio-morphological information on the distribution of SARS-CoV-2 infection foci obtained from LSFM analysis, high-resolution CLSM image stacks of a SARS-CoV-2 infection focus in the *Concha nasalis dorsalis* (ROI 1 from Figure 2) were acquired (Figure 5). Individual SARS-CoV-2-infected cells could be resolved, demonstrating cytoplasmic SARS-CoV-2 N distribution in both ciliated and non-ciliated cells (Figure 5B, arrows). Notably, SARS-CoV-2 N accumulated particularly at the apical side of the ciliated cells. Overall, these data demonstrate the feasibility of this correlated approach to dissect cell-specific responses to SARS-CoV-2 infection *in vivo* at subcellular resolution.

**Figure 5:**
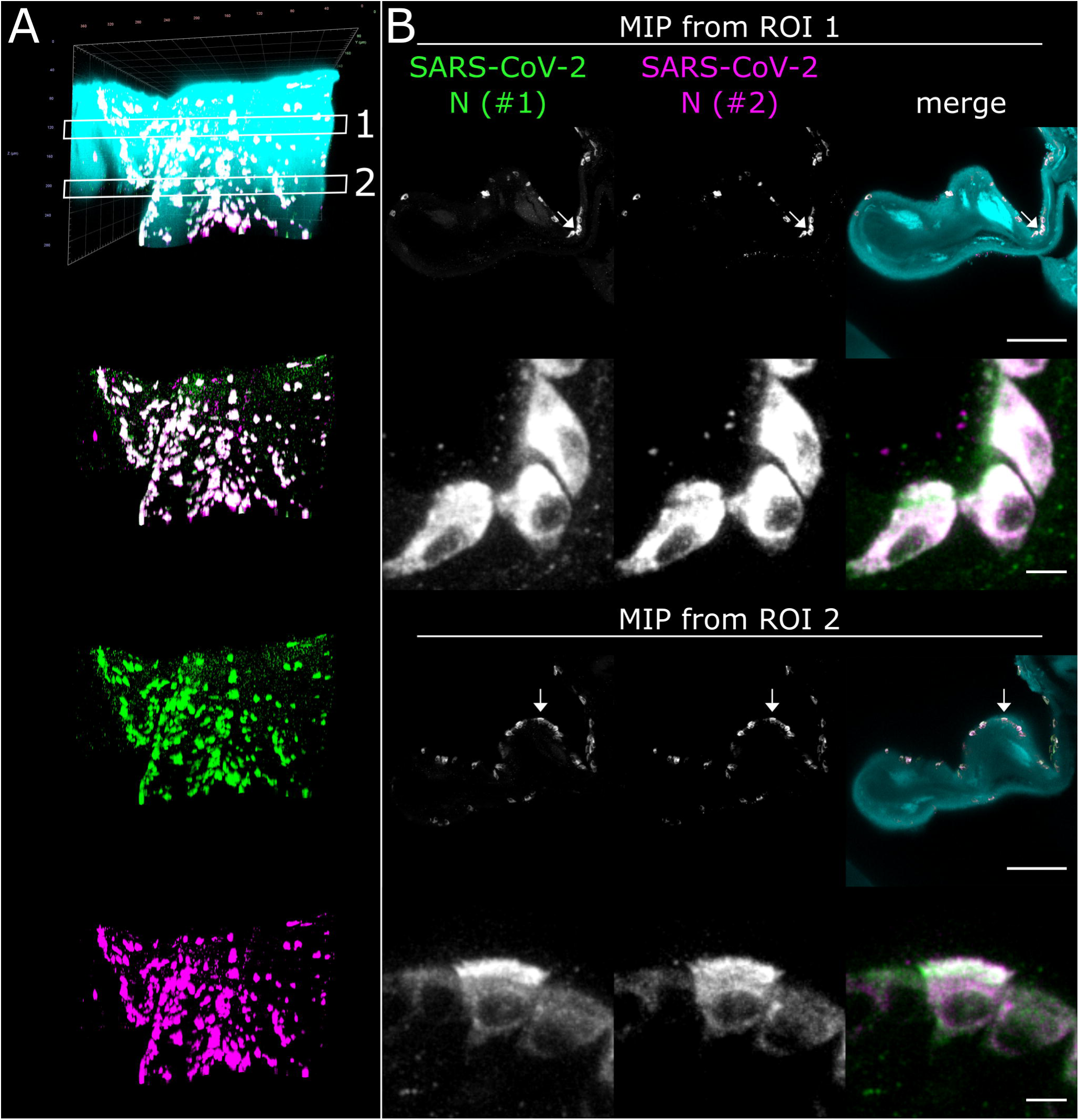
High-resolution CLSM analysis of SARS-CoV-2 infection foci in the concha nasalis dorsalis of a SARS-CoV-2-infected ferret at 4 days-post infection. **(A)** 3D maximum intensity projection (MIP) of a SARS-CoV-2 infection focus from ROI 1 in Figure 2. The image stack was acquired with a 40x/1.1 water immersion objective. Cyan = autofluorescence; green = SARS-CoV-2 N #1; magenta = SARS-CoV-2 N #2. Edge length of grid square = 40 µm. **(B)** MIPs from ROIs 1 and 2 in (A). Individual cells can be analyzed at subcellular resolution, highlighting infection of ciliated and non-ciliated cells (arrows). Scale bar = 100 µm (overview) and 5 µm (detail).

### SARS-CoV-2 detection in the LRT of ferrets

Previous studies demonstrated a preferential replication of SARS-CoV-2 in the URT of ferrets [29–31]. To assess whether comprehensive LSFM analysis may uncover previously undetected SARS-CoV-2 infection foci in the LRT, we looked at optically cleared high-volume lung and tracheal samples.

As before, some unspecific fluorescence signals could be seen in both lung (Figure 6) and tracheal tissue (Figure S2 and Supplementary Movie S3). At a first glance, no specific SARS-CoV-2 infection foci could be identified. However, hidden within an airway of the large lung tissue volume, a 172 µm by 102 µm-sized spot of colocalized antibody signals was detected (Figure 6B and Supplementary Movie S4). Contrary to the SARS-CoV-2 infection foci in the ferret nasal turbinates (Figures 2 and 3), the signal was localized above the epithelial cell layer (Figure 6B, single plane). This suggested detection of debris-associated antigen, which was mostly likely inhaled from the URT. Overall, while we were able to detect an 8.6×10^-5^ mm^3^ (86,000 µm^3^) spot of debris-associated antigen within a > 80 mm^3^ volume, we did not identify additional sites of infection within the LRT of ferrets, which corroborates preferential replication of SARS-CoV-2 in the URT of ferrets.

**Figure 6:**
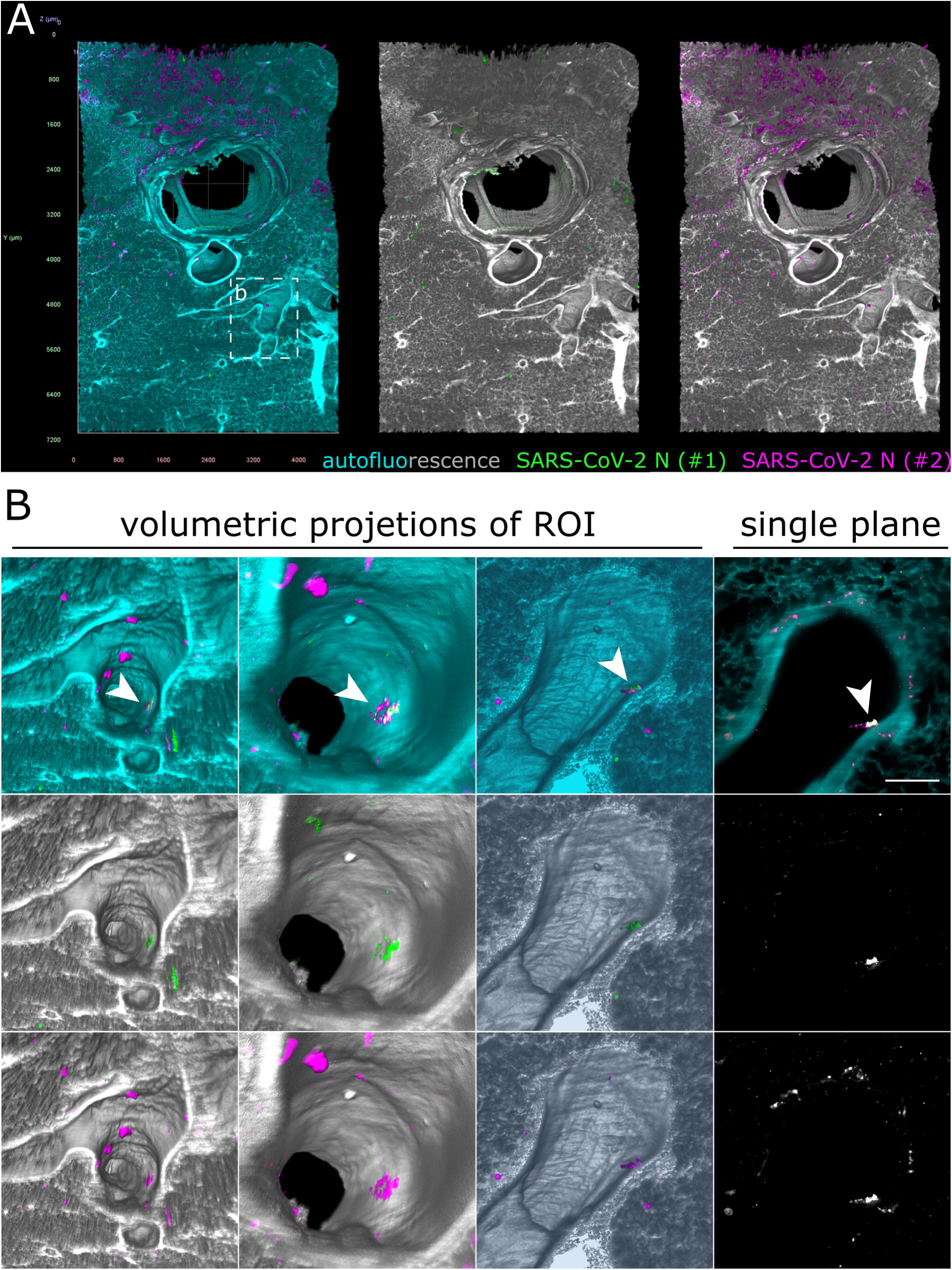
Only debris-associated SARS-CoV-2 antigen was detectable in ferret lung tissue at 4 days post-infection. **(A)** Volumetric projection of a large lung tissue section. While some background staining is detectable for the monoclonal antibody mix (#2, magenta), no signal overlap with the polyclonal antibody (#1, green) is visible. Cyan/grayscale = autofluorescence. Edge length of grid squares = 800 µm. Total magnification = 1.6x. **(B)** Alternate viewing angles reveal a spot inside an airway where both signals colocalize (white box in (A)). Contrary to the SARS-CoV-2-associated foci in Figures 2 and 3, the overlapping signal is detected lying on top the epithelial layer, suggesting that it is most likely cell debris inhaled from the URT.

## Discussion

While conventional immunohistochemistry studies have been used to assess the presence or absence of SARS-CoV-2 antigen in human and animal tissues [29–33,42,46,47], none of them were able to provide a greater spatial context of the infection site. By combining TOC with LSFM, we acquired large intact volumes of SARS-CoV-2-infected respiratory tissues from ferrets (Figure 1). The direct 3D visualization of virus infection via SARS-CoV-2 N staining established a comprehensive and mesoscopic overview of the infection in its full spatial context (Figures 2–6 and Supplementary Movies 1-4). Moreover, the determination of morphological parameters, e.g. focus-to-focus distances or the area of virus-infected tissue, not only allowed the characterization of individual SARS-CoV-2 infection foci but also provided a first quantitative insight into virus distribution within the spatio-morphological context of ferret nasal turbinates (Figure 4 and Table 1).

Here, we employed an ECi-based TOC approach [59] and adjusted it to visualize immunostained SARS-CoV-2 infection in large tissue samples of the respiratory tract of ferrets. While two 3D imaging approaches to SARS-CoV-2 infection in lung tissue have been reported, they have an entirely different scope: as both represent virtual histopathology strategies, they are meant to assess pathophysiology and associated tissue damage, but inherently cannot map and visualize specific SARS-CoV-2 infection. For the first study, Eckermann et al. [55] developed and demonstrated the utility of a phase contrast x-ray tomography concept to investigate unstained lung tissue. The second study describes the use of fluorescent H&E-analog stains (TO-PRO-3 for nuclear contrast and Eosin-Y for cytoplasmic/stromal contrast) to achieve “3D pseudo-histological imaging” [56]. Consequently, our study constitutes the first report of direct 3D visualization of SARS-CoV-2 infection via LSFM. While unspecific antibody signals were detected for both the polyclonal serum and the monoclonal mix directed against N of SARS-CoV, particularly on the outer surface of the tissue blocks, specificity for SARS-CoV-2 was ensured via colocalization of either staining (Figures 2, 3, 5, and 6) and absence of colocalization in naïve animals (Figure S1). Further optimization of the immunostaining protocol or the availability of SARS-CoV-2-specific antibodies will likely aid in reduction of background staining and improvement of virus detection.

In addition to the specific 3D reconstruction of SARS-CoV-2 infection within its spatio-morphological environment, the implementation of quantitative image analysis following accurate quantification of interrelated 3D parameters (Figure 4 and Table 1) represents a pronounced advantage of 3D immunofluorescence imaging over conventional IHC. To achieve a somewhat comparable yet more artifact-prone 3D reconstruction from thin sections, exceedingly laborious and time-consuming image registration pipelines following serial thin sectioning are necessary [61]. For instance, the nasal turbinate section from Figures 2–4 alone would require processing of around 800 sections (at 5 µm thickness), making it de facto impossible with 2D IHC.

When compared to previous studies [29–31], the spatial visualization of SARS-CoV-2 in the ferret respiratory tract confirmed preferential infection of the URT (Figures 2–4). Furthermore, our data indicate and emphasize a distinct oligofocal infection pattern of SARS-CoV-2 within nasal turbinates (Figures 2–4). Within a > 200 mm^3^ section of nasal turbinate tissue, only four SARS-CoV-2 infection foci (with a combined volume of 5.17×10^−3^ mm^3^) were detected, three of which accumulated in the *Concha nasalis dorsalis* and exhibited a maximum linear distance of 1.3 mm to each other (Figure 4 and Table 1). It is important to note that tissues inevitably shrink during the fixation, dehydration, and clearing process. For the EtOH-ECi-based TOC approach used here, a 50% volume reduction, equaling to a change of about 20% in tissue diameters, has been determined [59]. The limited degree of infection is particularly interesting in view of the amounts of infectious virus and genome copies that can be isolated from the URT of ferrets [29–31] and other animal species, like Syrian hamsters [32–34] and rhesus macaques [42–44]. Clustering of SARS-CoV-2 infection foci in narrow areas of the URT might also have implications for the likelihood of isolation of infectious virus and the detection of viral RNA from nasal swabs in comparison to nasal washes from ferrets and possibly other animal models. Accordingly, a high degree of variation in viral copy numbers can be observed from nose or throat swabs in comparison to bronchoalveolar lavages from SARS-CoV-2-infected rhesus macaques [42]. However, because of the proof-of-principle character of our study and the limited availability of SARS-CoV-2-infected material, further studies have to corroborate the clustering and focal infection pattern of SARS-CoV-2 in the URT.

Alongside complex quantitative 3D image analysis, volumetric imaging has the potential to discover rare events, as demonstrated by the detection of cancer metastases in sentinel lymph nodes, which had not been found via conventional IHC [62]. While preferential replication of SARS-CoV-2 in the URT of ferrets has been demonstrated via viral RNA and antigen detection [29–31], Kim et al. also detected some SARS-CoV-2-positive cells in the LRT. This is in contrast to the two other ferret susceptibility studies, which detected either no [31] or only low amounts [30] of viral RNA in the LRT, but neither found any SARS-CoV-2 antigen at this location. It is conceivable that scarce LRT infection had been overlooked in previous 2D IHC studies because of the focal character of SARS-CoV-2 infection in the tissue or that the detected viral RNA originated from URT-derived aspirated material. Using this high-volume imaging approach, we aimed to screen the tissue for rare SARS-CoV-2 infection foci in the LRT. While we did not detect any infection spots in tracheal tissue (Figure S2), we did visualize an individual, only 86,000 µm3-sized SARS-CoV-2 N-positive structure inside a lung airway (Figure 6B). However, spatial analysis revealed that the signal, contrary to the SARS-CoV-2 infection foci in the nasal turbinate epithelium (Figures 2 and 3), was detected above the airway epithelial layer. This strongly suggested that the structure most likely represents aspirated virus-containing debris as the result of localized cell or tissue damage at infected URT sites. Accordingly, this example emphasizes the suitability of this volumetric 3D LSFM approach to identify rare and highly localized pathogen-related events.

Ferrets are a standard model for human respiratory infection (reviewed in [63]). However, they recapitulate only mild SARS-CoV-2 infection and do not develop severe respiratory disease [11,29–31]. In contrast, both URT and LRT are strongly affected by SARS-CoV-2 infection in Syrian hamsters, including overt signs of disease [32,33]. This is closer to human disease, where SARS-CoV-2 antigen is found in the URT and LRT, as corroborated by the NHP model rhesus macaques [42–44]. While this proof-of-principle study is focused on ferret samples, it may serve as blueprint for further analyses in other animal models and even human clinical samples.

Independent of sample origin, volumetric imaging of cleared samples enables discovery and detection of rare infection events, as demonstrated earlier. This facilitates the investigation of the involvement of other organs outside of the respiratory tract in SARS-CoV-2 infection. For instance, viral antigen has been detected in the intestine of ferrets [29], hamsters [32,33], and rhesus macaques [42]. While viral RNA has been detected in human clinical brain samples [64], infection of cells of the central nervous system could thus far only be demonstrated experimentally in 3D human brain organoids [65]. Except for genetically modified mice expressing human angiotensin-converting enzyme 2 (ACE2) [39], the presence of SARS-CoV-2 antigen in animal or human brain samples has yet to be shown. Previous studies with other viral pathogens demonstrated that volumetric 3D imaging using TOC and LSFM is a highly valuable tool to assess the comprehensive distribution of virus infection *in vivo* [49,52]. Additional immunostaining against tissue-specific cell markers may further facilitate the investigation of the global SARS-CoV-2 cell tropism in affected tissues. Combining and correlating this with high-resolution CLSM analysis of SARS-CoV-2 hot spots identified via LSFM has the potential to dissect subcellular infection processes of SARS-CoV-2 *in vivo* with unparalleled detail. These results form the basis for research on larger sample sizes of both respiratory and non-respiratory tissues from SARS-CoV-2 animal models and human clinical samples using volumetric 3D LSFM of immunostained and cleared tissues.

Overall, we demonstrate the proof of concept for the utility of volumetric 3D immunofluorescence of critically important respiratory pathogens like SARS-CoV-2 using TOC and LSFM. The ability to analyze interrelated morphological parameters, like inter-foci distances and SARS-CoV-2-affected areas, and to put them into global perspective to the spatial tissue morphology provides unprecedented insight into SARS-CoV-2 infection in the respiratory tract of ferrets. In the future, this approach will be a crucial tool to understand the mesoscopic scale of host-pathogen interactions of SARS-CoV-2 but also other respiratory and non-respiratory pathogens, including, for example, influenza A and henipaviruses.

## Author contributions

Conceptualization, L.M.Z. and S.F.; methodology, L.M.Z.; formal analysis, L.M.Z.; investigation, L.M.Z., D.S. and M.M.; resources, J.S., D.H., M.B. and A.B.; writing— original draft preparation, L.M.Z. and S.F.; writing—review and editing, L.M.Z., D.S., J.S., M.M., D.H., M.B., E.M.A., T.C.M., A.B. and S.F.; visualization, L.M.Z.; supervision, E.M.A. and S.F.; project administration, S.F.; funding acquisition, S.F and T.C.M.

## Acknowledgments

We thank Roman Wölfel (German Armed Forces Institute of Microbiology, Germany) for providing the SARS-CoV-2 isolate and Sven Reiche (Friedrich-Loeffler-Institut, Germany) for providing the SARS-CoV N monoclonal antibodies. We are grateful to Angela Hillner for excellent technical assistance. Figure 1A was created with Biorender.com.

