## Supplementary Materials (Supplementary Figures + Video Captions) for "3D reconstruction of SARS-CoV-2 infection in ferrets emphasizes focal infection pattern in the upper respiratory tract"

### Supplementary Material

**Supplementary Figure S1: No SARS-CoV-2 colocalization of antibody signals is observable in tissue from mock-infected control animals.** Volumetric projections of nasal turbinate (A), lung (B), and trachea (C) tissue from mock-infected control animals. No colocalization of the polyclonal serum (#1, green) and the monoclonal mix (#2, magenta) can be detected. Cyan/grayscale = autofluorescence. Edge length of grid squares = 1.000 µm. Total magnification = 1.6x (A and B), 2x (C).

**Supplementary Figure S2: No SARS-CoV-2 infection foci were detected in ferret tracheal tissue.** Volumetric projection of a large ferret trachea section. Only unspecific background staining is detectable for the polyclonal serum (#1, green) and the monoclonal antibody mix (#2, magenta). No signal overlap of both antibody signals was observable. Cyan/grayscale = autofluorescence. Edge length of grid squares = 800 µm. Total magnification = 2x.

**Supplementary Table S1: List of chemicals and reagents.**

**Supplementary Movie S1: Volumetric 3D projection of an LSFM-acquired, > 200 $mm^3$-sized nasal turbinate section from a SARS-CoV-2-infected ferret.** The tissue morphology was reconstructed using non-specific tissue autofluorescence (cyan). Edge length of grid squares = 2 mm. Total magnification = 1.26x.

**Supplementary Movie S2: Fly-through animation of individual SARS-CoV-2 infection foci in ferret nasal turbinates at 4 days post-infection.** The three distinct SARS-CoV-2 infection foci from Figures 3 and 4 are highlighted at timestamps (mm:ss) 00:09 [A1], 00:14 [A5], and 00:25 [A7]. Cyan = autofluorescence; green = SARS-CoV-2 N #1; magenta = SARS-CoV-2 N #2. Edge length of grid squares = 300 µm. Total magnification = 8x.

**Supplementary Movie S3: 360° rotation of a trachea section from a SARS-CoV-2-infected ferret at 4 days post-infection.** No SARS-CoV-2-associated infection spots were detected. Cyan = autofluorescence; green = SARS-CoV-2 N #1; magenta = SARS-CoV-2 N #2. Edge length of grid squares = 800 µm. Total magnification = 2x.

**Supplementary Movie S4: Fly-through animation of likely debris-associated SARS-CoV-2 infection in ferret lung tissue.** The colocalization of either SARS-CoV-2 N antibody signal (#1, green; #2, magenta) inside a lung airway can be observed at timestamp (mm:ss) 00:16. Cyan = autofluorescence. Edge length of grid squares = 200 µm. Total magnification = 6.4x.

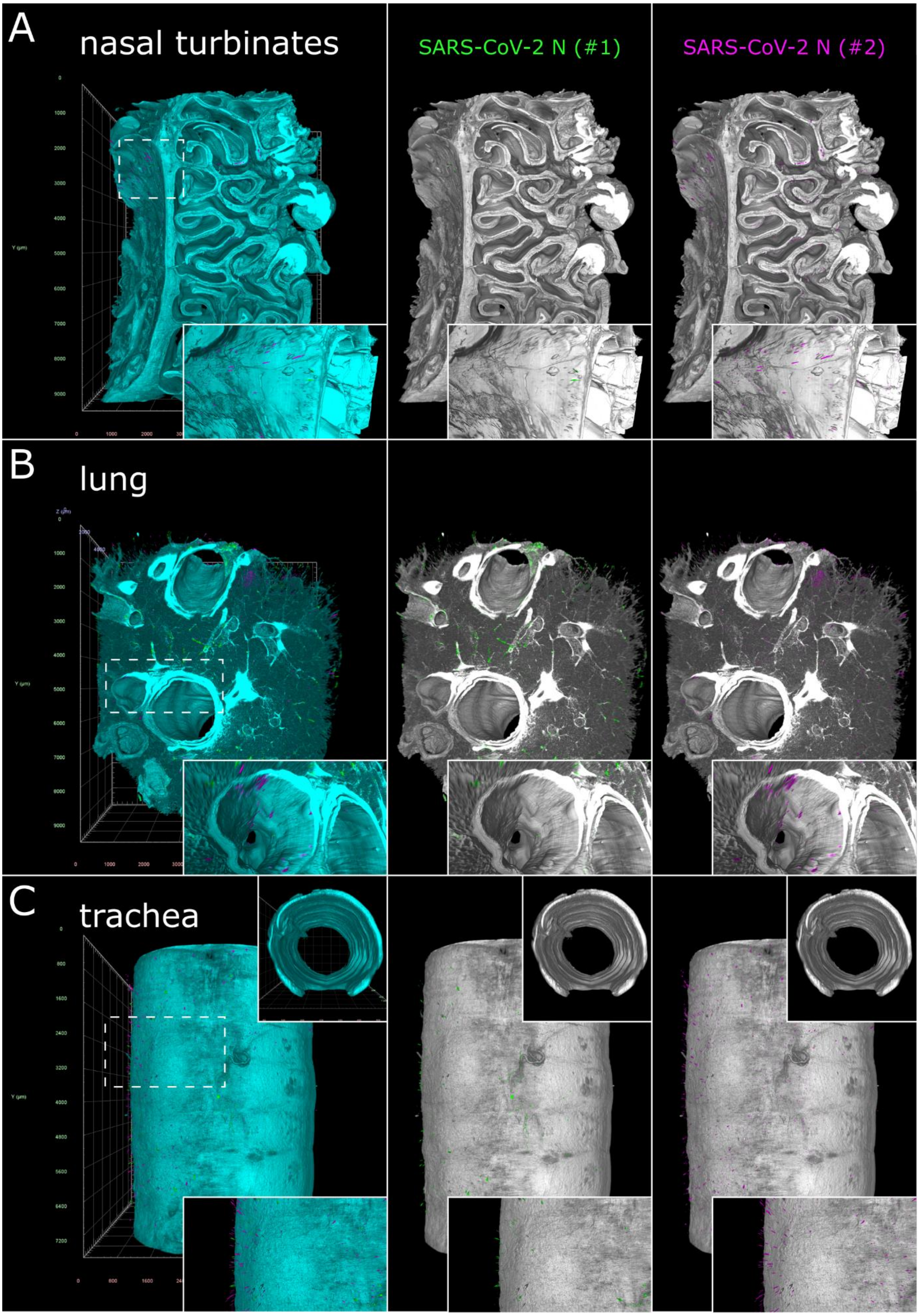


**Supplementary Figure S1: No SARS-CoV-2 colocalization of antibody signals is observable in tissue from mock-infected control animals.** Volumetric projections of nasal turbinate **(A)**, lung **(B)**, and trachea **(C)** tissue from mock-infected control animals. No colocalization of the polyclonal serum (#1, green) and the monoclonal mix (#2, magenta) can be detected. Cyan/grayscale = autofluorescence. Edge length of grid squares = 1,000 µm. Total magnification = 1.6x (A and B), 2x (C).

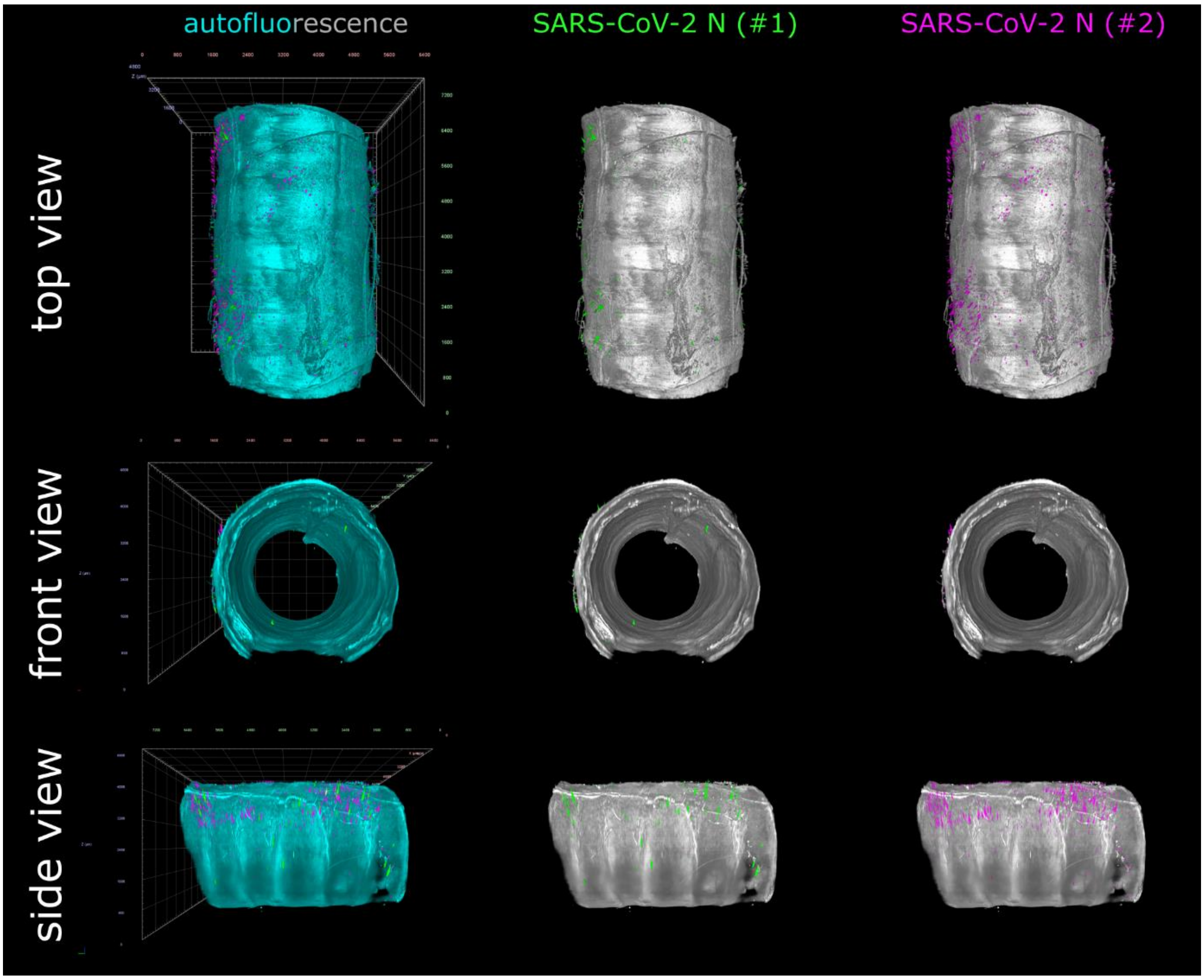


**Supplementary Figure S2: No SARS-CoV-2 infection foci were detected in ferret tracheal tissue at 4 days post-infection.** Volumetric projection of a large ferret trachea section. Only unspecific background staining is detectable for the polyclonal serum (#1, green) and the monoclonal antibody mix (#2, magenta). No signal overlap of both antibody signals was observable. Cyan/grayscale = autofluorescence. Edge length of grid squares = 800 µm. Total magnification = 2x.

**Supplementary Table 1: List of chemicals and reagents.**

| reagents | source | PO number | additional information |
|---|---|---:|---|
| DMSO | Carl Roth | 4720 | dimethyl sulfoxide |
| ethanol | Carl Roth | 9065 | dehydrating agent |
| ethyl cinnamate | Alfa Aesar | A12906 | clearing agent |
| Formical-2000™ | Statlab | 1314 | decalcifier |
| glycine | Carl Roth | 3908 | autofluorescence quencher |
| heparin sodium salt | Carl Roth | 7692 | reduction of background |
| hydrogen peroxide | Carl Roth | 8070 | bleaching agent |
| n-hexane | Alfa Aesar | 43263 | delipidating agent |
| normal donkey serum | Bio-Rad | C06SBZ | blocking agent |
| Triton X-100 | Carl Roth | 3051 | detergent |
| Tween-20 | AppliChem | A4974 | detergent |
